# Is there a need for preoperative α-blocker in patients missed preoperative diagnosis of extra-adrenal retroperitoneal paraganglioma undergoing paraganglioma resection? A retrospective study of 167 cases at a single center

**DOI:** 10.1101/2020.09.21.305870

**Authors:** Yi Liu, Xinye Jin, Jie Gao, Shan Jiang, Lei Liu, Jing-Sheng Lou, Bo Wang, Hong Zhang, Qiang Fu

**Affiliations:** Department of Anesthesiology, the first Medical Center of Chinese PLA General Hospital, Beijing 100853, China; Department of Anesthesiology, the third Medical Center of Chinese PLA General Hospital, Beijing, 100039; Department of Endocrinology, Hainan Hospital of PLA General Hospital, Sanya 572013, Hainan Province, China; Department of Endocrinology, the first Medical Center of Chinese PLA General Hospital, Beijing 100853, China; Department of Anesthesiology, Tsinghua University Affiliated Beijing Tsinghua Changgung Hospital, Beijing 102218, China

**Keywords:** Paraganglioma, Premedication, Anesthesia, Adrenergic Alpha-Antagonists, Hemodynamics, Outcomes

## Abstract

**Background:** Preoperative α-adrenergic blockade is believed to decrease perioperative risks and mortality in adrenal pheochromocytoma surgeries. The aim of this study is to evaluate the effects of the preoperative α-adrenergic blockade on patients’ outcomes in extra-adrenal retroperitoneal paraganglioma surgeries.

**Methods:** We searched our clinical database for the diagnosis extra-adrenal retroperitoneal paraganglioma by postoperative histopathology in the General Hospital of People’s Liberation Army from 2000 till 2017. And we recorded preoperative status of patients, preoperative medication preparation, intraoperative and postoperative cardiovascular events, intake and output, length of stay in ICU, length of hospital stay, and short time outcomes.

**Results:** The intraoperative morbidity of heart rate elevation and highest heart rate were higher in patients undergoing tumor manipulation with preoperative α-adrenergic blockade than those without (*P*<0.05), while there were no significant differences in intraoperative morbidity of blood pressure elevation and SAP decreased following tumorectomy in these two groups (*P*>0.05). There were no significant differences in postoperative complications and outcomes (*P*>0.05).

**Conclusion:** Under the current medical techniques, either with or without preoperative medicine, resection of extra-adrenal retroperitoneal paraganglioma could be carried out successfully.

## Introduction

Pheochromocytoma removal surgery is a risky procedure involving high perioperative morbidity and mortality due to hemodynamic instability. Recent studies showed that the decline of perioperative mortality associated with pheochromocytoma resection from 20%-45% to 0%-2.9% was attributed to the development of imaging techniques, surgical and anesthetic techniques and preoperative medical management[1, 2]. Preoperative α-adrenergic blockade, in particular, is believed to be the major factor to reduce the risk of intraoperative hemodynamic instabilities despite the absence of randomized controlled trials[3, 4]. Theoretically, preoperative management should strictly follow these criteria to decrease systemic vascular resistance, increase venous compliance, expand volume and reduce the risk of hypovolemic shock after tumor removal[4-6]. Many retrospective studies showed operations with preoperative preparation could have better outcomes than those without preoperative agents.

But most extra-adrenal paraganglioma patients, with no accompanying clinical symptoms such as hypertension, headache, or palpitation and lacking of functional imaging, could be misdiagnosed as other types of retroperitoneal masses. There were a few case reports about such operations with negligible complications intraoperatively and postoperatively, while no large-scale sample assessment was conducted[7-9]. Under such circumstances, the aim of the present retrospective study was to verify whether the patients with extra-adrenal retroperitoneal paraganglioma have bad outcomes when they were not accompanied with preoperative α-adrenergic blockade.

## Materials and methods

### Patients

With the approval of the Chinese PLA General Research Ethics Committee, we performed a unicentral retrospective analysis of patients diagnosis as retroperitoneal paraganglioma. We screened our administrative diagnosis database for patients with retroperitoneal neoplasm (International Classification of Diseases (ICD)-10 code C48.0, C75.7, C78.6, D20.0 and D48.3; n = 4457). Extra-adrenal retroperitoneal paraganglioma was defined according to the postoperative pathological examination. For patients who had repeated operations, only the first surgical procedure was included. Thus, we identified 167 patients with retroperitoneal paraganglioma that met the inclusion criteria. The retrospective study followed the tenets of the Declaration of Helsinki for research involving human subjects. In accordance with Chinese law, retrospective studies on medical records performed in China do not need written consent from participants.

### Perioperative parameter

Operation duration was measured from the cutting of skin to the end of incision. Intraoperative hemodynamic instability (HI) was defined as follows: systolic blood pressure (SBP)over 180 mmHg more than 10 minutes was designated as cut-off value for intraoperative hypertension and SBP below 80 mmHg as hypotension; heart rate (HR) over 120 beat per minute (bpm) more than 10 minutes was regarded as tachycardia and HR below 50 bpm as bradycardia[10, 11].

Preoperative, intraoperative and postoperative data were collected and compared between patients with or without preoperative α-blockage.

Preoperative data: age, sex, weight, preoperative blood pressure, preoperative heart rate, accompanying disease, symptoms, definite diagnosis or not and preoperative ECG abnormal or not.

Intraoperative data: duration of surgery, intraoperative cardiovascular events, output and input, administration of cardioactive drugs, major axis of tumor and operation approach.

Postoperative data: postoperative stay, ICU stay and postoperative short-term outcomes.

### Statistical analysis

Statistical analysis was performed using IBM SPSS Statistics 22 (SPSS Inc, Chicago, Illinois, USA). Normally distributed data were presented as mean ± standard deviation. Non-normally distributed data were presented as median (interquartile). The Student t-test, the Mann-Whitney u-test, ANOVA and the Chi-square test were employed as required. We looked for univariate associations between the presence or absence of unstable intraoperative hemodynamics, including intraoperative hypertension or intraoperative hypotension, surgical approach, the following patient and tumor characteristics: age, sex, history of cardiovascular events, the presence of an underlying familial disease, preoperative BP levels, preoperative HR level, with preoperaive αblocke or not, preoperative complications, preoperative abnormal electrocardiogram, tumor diameter. *P* < 0.05 were considered significant difference.

## Results

### Patient characteristics

Patient characteristics are shown in Table 1. Sixty-eight patients were diagnosed with extra-adrenal retroperitoneal paraganglioma preoperatively. Sixty-one of these patients with definite diagnosis were pretreated with α-adrenergic receptor antagonists at least for 2 weeks, including phenoxybenzamine, phentolamine, or terazosin, in order to normalize the preoperative blood pressure and HR. Seven patients diagnosed as extra-adrenal retroperitoneal paraganglioma did not receive α-blockers because of normal blood pressure preoperatively, and the other 99 patients without preoperative definite diagnosis underwent operations for the removal of occupying lesions. The patients with preoperative α-blockers suffered higher incidence rates of preoperative hypertension and diabetes, and revealed a higher incidence of headache, dizzy, sweat, chest stress, palpitation and elevation of blood pressure and blood glucose (*P* < 0.05). After preoperative preparation, the preoperative SAP and HR in the two groups were similar (*P* > 0.05).

**Table 1.**
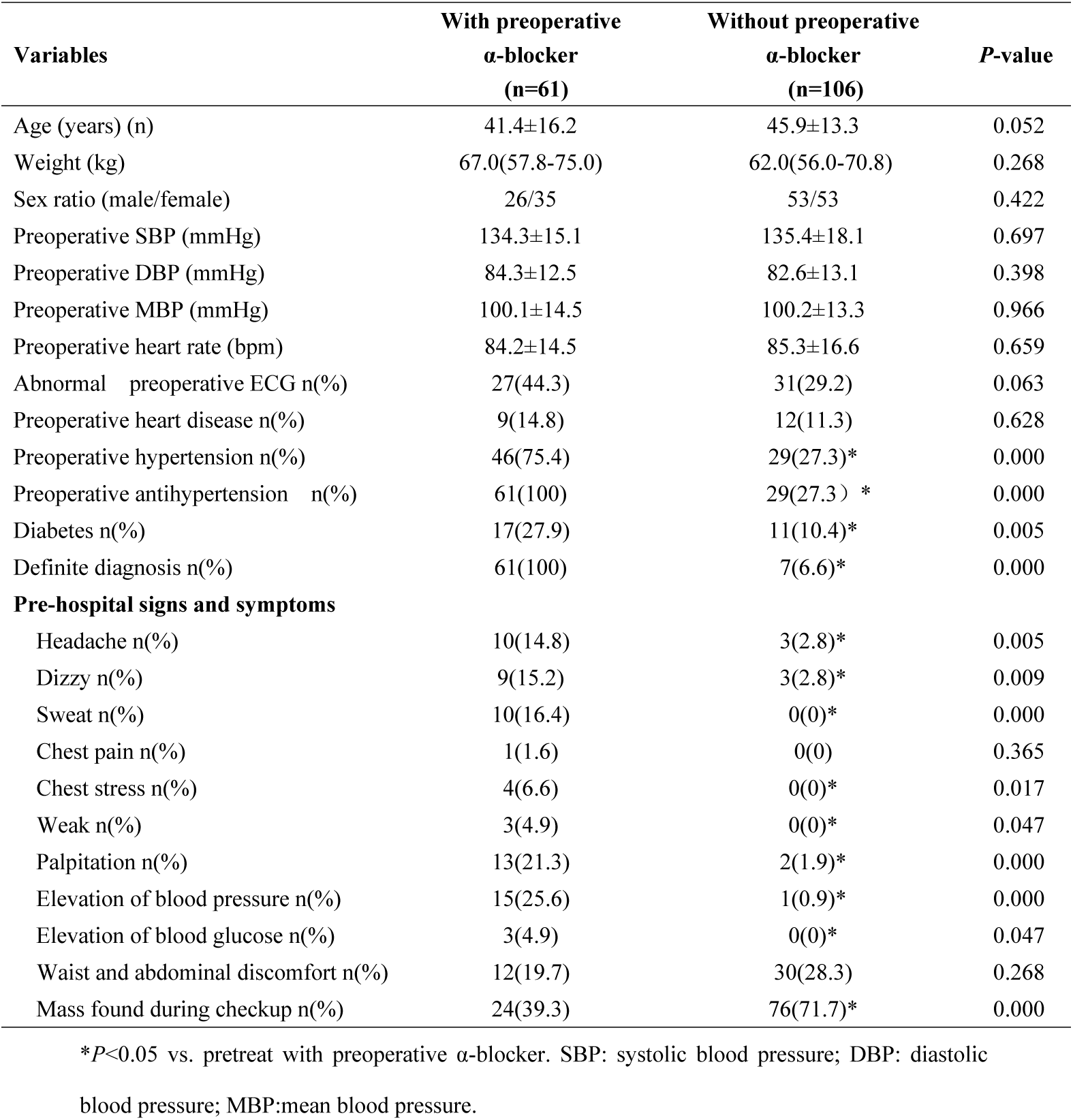
Patient characteristic

| Variables | With preoperative<br>α-blocker<br>(n=61) | Without preoperative<br>α-blocker<br>(n=106) | P-value |
| --- | --- | --- | --- |
| Age (years) (n) | 41.4±16.2 | 45.9±13.3 | 0.052 |
| Weight (kg) | 67.0(57.8-75.0) | 62.0(56.0-70.8) | 0.268 |
| Sex ratio (male/female) | 26/35 | 53/53 | 0.422 |
| Preoperative SBP (mmHg) | 134.3±15.1 | 135.4±18.1 | 0.697 |
| Preoperative DBP (mmHg) | 84.3±12.5 | 82.6±13.1 | 0.398 |
| Preoperative MBP (mmHg) | 100.1±14.5 | 100.2±13.3 | 0.966 |
| Preoperative heart rate (bpm) | 84.2±14.5 | 85.3±16.6 | 0.659 |
| Abnormal preoperative ECG n(%) | 27(44.3) | 31(29.2) | 0.063 |
| Preoperative heart disease n(%) | 9(14.8) | 12(11.3) | 0.628 |
| Preoperative hypertension n(%) | 46(75.4) | 29(27.3)* | 0.000 |
| Preoperative antihypertension n(%) | 61(100) | 29(27.3) * | 0.000 |
| Diabetes n(%) | 17(27.9) | 11(10.4)* | 0.005 |
| Definite diagnosis n(%) | 61(100) | 7(6.6)* | 0.000 |
| **Pre-hospital signs and symptoms** |  |  |  |
| Headache n(%) | 10(14.8) | 3(2.8)* | 0.005 |
| Dizzy n(%) | 9(15.2) | 3(2.8)* | 0.009 |
| Sweat n(%) | 10(16.4) | 0(0)* | 0.000 |
| Chest pain n(%) | 1(1.6) | 0(0) | 0.365 |
| Chest stress n(%) | 4(6.6) | 0(0)* | 0.017 |
| Weak n(%) | 3(4.9) | 0(0)* | 0.047 |
| Palpitation n(%) | 13(21.3) | 2(1.9)* | 0.000 |
| Elevation of blood pressure n(%) | 15(25.6) | 1(0.9)* | 0.000 |
| Elevation of blood glucose n(%) | 3(4.9) | 0(0)* | 0.047 |
| Waist and abdominal discomfort n(%) | 12(19.7) | 30(28.3) | 0.268 |
| Mass found during checkup n(%) | 24(39.3) | 76(71.7)* | 0.000 |

\* $P < 0.05$ vs. pretreat with preoperative α-blocker. SBP: systolic blood pressure; DBP: diastolic
blood pressure; MBP:mean blood pressure.

### Intraoperative parameters

Compared with patients without preoperative α-blockers, those administered with preoperative α-blocker had higher intraoperative HR (114.57±22.03 bpm vs. 104.60±17.58 bpm) and higher morbidity of HR (42.6% vs. 22.6%) elevation during tumor manipulation (*P* < 0.05) (Table 2). There were no differences in surgery duration, morbidities of intraoperative SAP elevation when tumor manipulation, SAP decreased following tumorectomy, fluid intake and bleeding (*P* > 0.05) (Table 2). The tumor diameters in the patients without preoperative α-blockers were larger than those without α-blockers (6.0 (3.0) cm vs. 5.5 (3.0) cm, *P* < 0.05). The laparoscope or robot was more used in the patients pretreated with α-blockers (*P* < 0.05) (Table 2). Patients undergoing preoperative α-blockers were more likely to take intraoperative anti-hypertensive drugs, particularly phentolamine, esmolol and sodium nitroprusside to normalize the intraoperative hemodynamics (*P* < 0.05). (Figure 1)

**Table 2.** Surgery duration, intraoperative cardiovascular events, intake and output, tumor diameters and operation approach.

| Variables | With preoperative<br>α-blocker<br>(n=61) | Without preoperative<br>α-blocker<br>(n=106) | P-value |
| --- | --- | --- | --- |
| Duration of surgery (min) | 178.69±79.88 | 187.85±87.36 | 0.503 |
| Highest intra-operative SAP<br>before tumor resection (mmHg) | 179.98±39.12 | 173.53±36.83 | 0.288 |
| SAP>180mmHg n(%) | 28(45.9) | 45(42.5) | 0.746 |
| Lowest intra-operative SAP<br>after tumor resection (mmHg) | 91.84±21.60 | 91.36±15.84 | 0.870 |
| SAP<80mmHg n(%) | 15(26.2) | 26(24.5) | 1.000 |
| Highest intra-operative heart rate before<br>tumor resection (bpm) | 114.57±22.03 | 104.60±17.58* | 0.002 |
| Heart rate>120 bpm n(%) | 26(42.6) | 24(22.6)* | 0.009 |
| Bleeding (ml) | 300 (850) | 300 (700) | 0.499 |
| Urine (ml) | 650(600) | 500(635) | 0.235 |
| Crystalloid (ml) | 2550(1635) | 2100(1400) | 0.327 |
| Colloid (ml) | 1000(1000) | 1250 (500) | 0.900 |
| Red cell (ml) | 0 (400) | 0(600) | 0.655 |
| Plasm (ml) | 0 (225) | 0(225) | 0.501 |
| Major axis of tumor (cm) | 5.5 (3.0) | 6.0 (3.0)* | 0.042 |
| Operation approach: laparoscope or robot<br>n(%) | 21 (34.4) | 9 (5.7)* | 0.000 |

\* $P<0.05$ vs. pretreat with preoperative α-blocker. SBP: systolic blood pressure; DBP: diastolic blood pressure; MBP:mean blood pressure.

**Figure 1.**
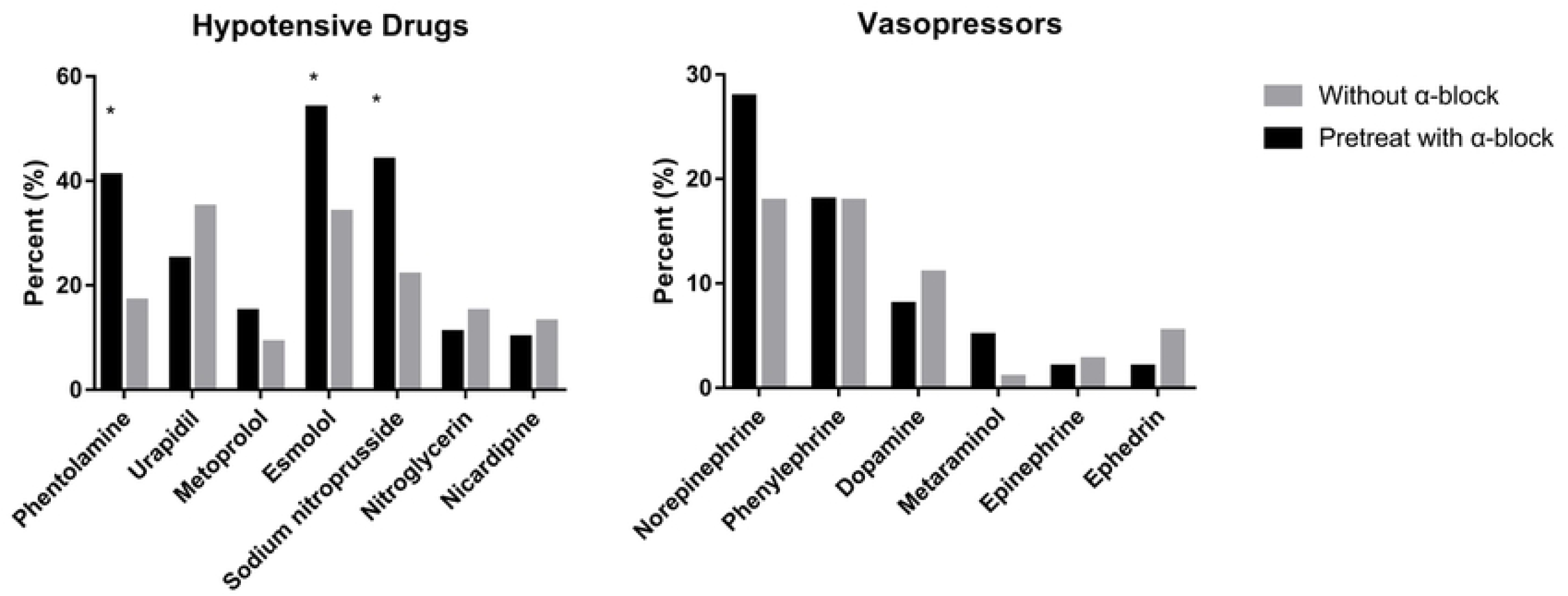
The intraoperative employment of vasoactive drugs. \**P*<0.05 vs. pretreat without preoperative α-blocker.

### Surgical outcomes

As shown in Table 3, there were no differences in the total duration of hospital stay, postoperative stay and ICU stay (*P* > 0.05). However, the patients pretreated with α-blockers stayed longer at the hospital before operation than those without preoperative α-blockers (10.0(12.0) days vs. 6.5(5) days, *P* < 0.05). One patient without preoperative α-blockers died the first day after surgery due to hemorrhagic shock in the ICU after uncontrolled intraoperative massive hemorrhage. Another patient without preoperative α-blockers died the third day after surgery for regurgitation and aspiration. There were no differences in postoperative complications and outcomes between the two groups.

**Table 3.**
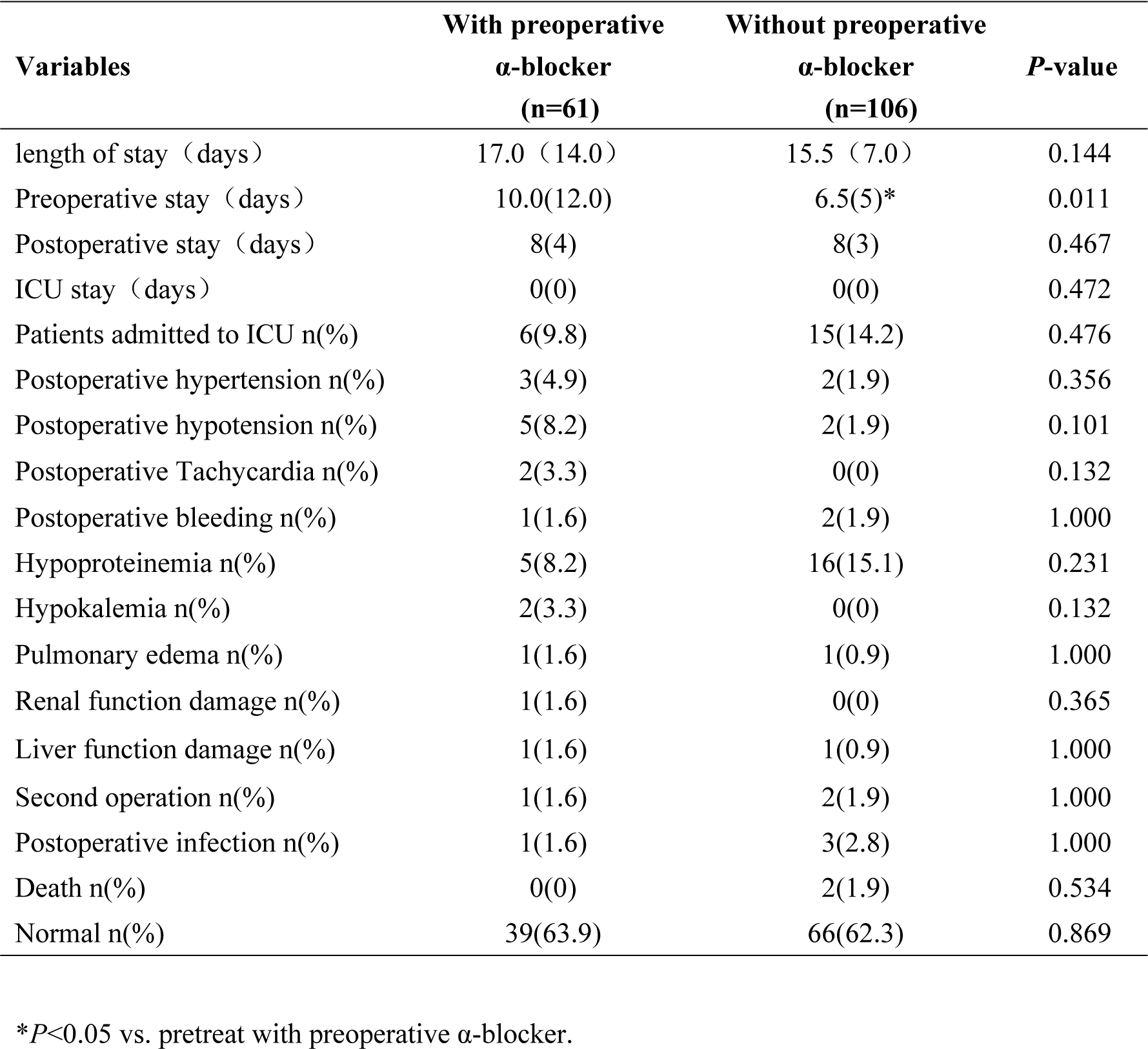
Postoperative complications and short time outcomes

| **Variables** | **With preoperative**<br>**α-blocker**<br>**(n=61)** | **Without preoperative**<br>**α-blocker**<br>**(n=106)** | **P-value** |
| --- | --- | --- | --- |
| length of stay (days) | 17.0 (14.0) | 15.5 (7.0) | 0.144 |
| Preoperative stay (days) | 10.0(12.0) | 6.5(5)* | 0.011 |
| Postoperative stay (days) | 8(4) | 8(3) | 0.467 |
| ICU stay (days) | 0(0) | 0(0) | 0.472 |
| Patients admitted to ICU n(%) | 6(9.8) | 15(14.2) | 0.476 |
| Postoperative hypertension n(%) | 3(4.9) | 2(1.9) | 0.356 |
| Postoperative hypotension n(%) | 5(8.2) | 2(1.9) | 0.101 |
| Postoperative Tachycardia n(%) | 2(3.3) | 0(0) | 0.132 |
| Postoperative bleeding n(%) | 1(1.6) | 2(1.9) | 1.000 |
| Hypoproteinemia n(%) | 5(8.2) | 16(15.1) | 0.231 |
| Hypokalemia n(%) | 2(3.3) | 0(0) | 0.132 |
| Pulmonary edema n(%) | 1(1.6) | 1(0.9) | 1.000 |
| Renal function damage n(%) | 1(1.6) | 0(0) | 0.365 |
| Liver function damage n(%) | 1(1.6) | 1(0.9) | 1.000 |
| Second operation n(%) | 1(1.6) | 2(1.9) | 1.000 |
| Postoperative infection n(%) | 1(1.6) | 3(2.8) | 1.000 |
| Death n(%) | 0(0) | 2(1.9) | 0.534 |
| Normal n(%) | 39(63.9) | 66(62.3) | 0.869 |

\* $P < 0.05$ vs. pretreat with preoperative α-blocker.

### Subgroup analysis

When compared with patients without preoperative α-adrenergic blockade, the patients taking other preoperative hypotensors and without any preoperative hypotensive drugs experienced the same intraoperative and postoperative circulation changing and had the same outcomes (Table 4).

**Table 4.**
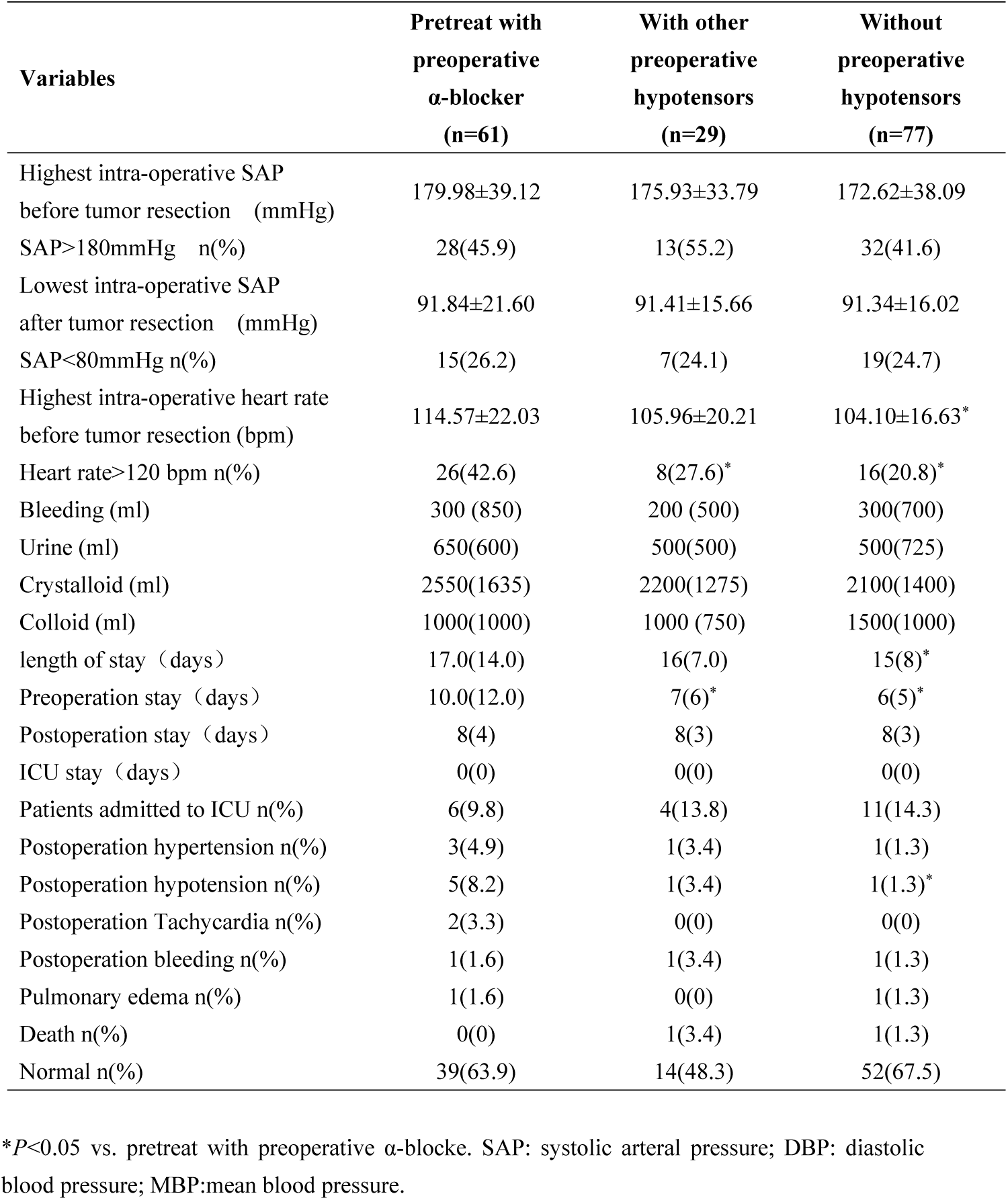
Intraoperative cardiovascular events, Postoperative complications and short time outcomes of subgroup analysis.

## Discussion

This is the first large-scale retrospective study of patients with extra-adrenal retroperitoneal paraganglioma, most of whom were undiagnosed properatively. And our research revealed that paraganglioma resection could be carried out successfully without preoperative α-blocker in patients of omission diagnosis of paraganglioma preoperatively.

Serials of studies show that, about 80% to 85% of pheochromocytoma are located in adrenal medulla, which are called chromaffinoma or pheochromocytoma, whilst 15% to 20% are extra-adrenal, which are called paraganglioma and usually located close to the sympathetic chain, such as in the head and neck, thoracic cavity, and retroperitoneal cavity[11-13]. Extra-adrenal retroperitoneal paraganglioma, with no prominent clinical manifestations like headache, perspiration, and palpitations resulting from the release of catecholamine, was prone to be misdiagnosed as other retroperitoneal masses [14-16]. Compared with adrenal pheochromocytoma, extra-adrenal retroperitoneal paraganglioma has some significant characteristics, such as high misdiagnosis rate, complicated anatomic structure, etc [17, 18]. Regarding the characteristics of extra-adrenal retroperitoneal paraganglioma, most of our patients had not manifested typical clinical symptoms of catecholamine release, so that surgeons had not been aware of extra-adrenal retroperitoneal paraganglioma which resulted in most of our patients having no preoperative medicines.

Perioperative hemodynamic instability was believed to increase the perioperative mortality and morbidity [19]. Lacking evidence from randomized controlled clinical studies, a lot of retrospective studies and institutional experience suggested paraganglioma patients must take preoperative α-blocker in order to reduce perioperative hemodynamic instability [20-23]. Conversely, some studies showed patients would undergo safe surgical procedure without preoperative α-blocking agents [7, 24]. Boutros *et al*. reported that all the 29 patients in their series without using preoperative α-adrenergic blockade survived and were discharged from hospital without clinical evidence of cardiovascular complications and proved that patients with pheochromocytoma could undergo successful surgery without preoperative profound and long-lasting alpha adrenergic blockade. All their patients were confirmed preoperatively and infused with sodium nitroprusside and nitroglycerin alone or in combination intraoperatively [25]. Similarly, Lentchener *et al*. reported that high preoperative SAP was not indicative of intra- and postoperative hemodynamic instability with no regard to the administration of preoperative hypotensive drugs [26]. In our study, 29 patents who were suffered from hypertension in the group without preoperative α-adrenergic blockade took β-blockers, calcium channel blocker and angiotensin-converting enzyme inhibitors to normalize the blood pressure, like metoprolol, nifedipine, nimodipine and captopril and so forth, alone or in combination. Under the subgroup analysis, the intraoperative and postoperative circumstances of the patients with antihypertensive drugs in group without preoperative α-blocker were same when compared with the patients with preoperative α-blocker. It seemed the other kind types of hypotensors not only α-adrenergic blockade could be used safely as the preoperative medicine for extra-adrenal retroperitoneal paraganglioma. The patients with α-blocker had prolonged hospital stay especial preoperative stay for normalizing the preoperative blood pressure according to the routine recommendation to take preoperative α-blocker at least for 2 weeks, and could induce intraoperative tachycardia.

All paraganglioma were believed to synthesize and store catecholamine, and functional tumors were defined as having elevated urine or serum catecholamine levels attributed to the presence of tumor [27]. Although our study had some limitations in that none of our patients had intraoperative blood serum catecholamine assay test, about 40% patients had experienced hemodynamic instabilities including elevated blood pressure and heart rate during tumor manipulation. Therefore we can only assume those tumors with intraoperative hemodynamic instabilities were functional paraganglioma. Tauzin *et al*. in their series showed that there was no correlation between preoperative urinary metanephrine and normetanephrine levels and intraoperative plasma catecholamine concentrations, but all their patients received preoperative α-adrenergic blockade at least for 15 days[3]. Intraoperative catecholamine release depends mainly on intubation, first incision, peritoneal insufflation, surgical manipulation of the tumor and tumor diameters, and these can result in dramatic intraoperative hemodynamic variations and huge challenges to anesthesiologists[3, 5, 28]. According to the hemodynamic changes in our study, it seemed to be no differences in the abilities of intraoperative catecholamine release in the two groups. For the shortage of retrospective study, we had no tests of intraoperative plasma catecholamine level, and we could design a prospective study to measure the plasma catecholamine in various time points intraoperatively.

Extra-adrenal retroperitoneal paraganglioma have close relationship with abdominal aorta, inferior vena cava, renal artery, renal vein and other retroperitoneal organs, and consequently, the operations should always be conducted by open approaches and could experience massive hemorrhage due to the complicated structure [29-31]. With the rapid development of surgical skills, more and more patients with pheochromocytoma derived from adrenal medulla undergo retroperitoneal laparoscopic approach [32-34]. The same goes in extra-adrenal retroperitoneal paraganglioma. In our study, 30 patients underwent operations with laparoscope or robot-assisted laparoscope from 2009. This approach was more likely to be applied in the patients with preoperative α-blocker because of small tumor diameters and definite diagnosis. Meanwhile, most of our patients experienced massive hemorrhage, and more than 27% of our patient had blood loss larger than 800 ml, and needed massive blood transfusion and fluid therapy. It seemed that preoperative α-blockers could not decrease the risk of massive bleeding in resection of retroperitoneal tumors.

Most of our patients had good outcomes without significant complications whether they received preoperative α-blocker or not, which could be attributed to the successful intraoperative and postoperative managements by surgeons and anesthesiologists [4, 28, 35]. Previous studies made the surgeons and anesthesiologists aware of the pathophysiology of paraganglioma, especially the dramatic changes of circulation due to the variations of both catecholamine release and the volume during intraoperative period [36]. Real-time dynamic circulation parameters could be displayed without delay from invasive monitor, anesthesiologists could deal with these events with plenty of fast-acting and short-term drugs, like sodium nitroprusside, urapidil, esmolol, nicardipine and norepinephrine[3, 36]. Recently, more and more new techniques are employed in surgery to carry out accurate evaluation of circulating blood volume and goal-directed volume therapy in order to improve outcomes and reduce hospital stay [37, 38]. Furthermore, some preoperative interventional therapies are employed in order to reduce the risk of intraoperative catecholamine release induced by tumor manipulation [39, 40].

There are several limitations in our study. Firstly, due to the high rate of preoperative misdiagnosis of extra-adrenal retroperitoneal paraganglioma, most of the cases did not have preoperative blood or urine catecholamine examinations. As well, there were no tests of intraoperative blood or urine catecholamine examinations. Secondly, this study included nearly 20 years of cases from 2000 to 2017, as surgical techniques developed from traditional open approach surgery to minimally invasive surgery like robot-assisted surgery. Nevertheless, there were no differences in the perioperative hemodynamic instabilities and postoperative outcomes no matter what kinds of surgery approaches were applied.

## Conclusion

In conclusion, our findings demonstrated that most of extra-adrenal retroperitoneal paraganglioma could experience intraoperative hemodynamic instabilities, whether preoperative α-blocker was given or not. Moreover, ensured by current surgical approaches, anesthesia skills, monitor technologies and cardiovascular drugs, patients who were missed preoperative diagnosis of extra-adrenal retroperitoneal paraganglioma could undergo surgery successfully and safely without preoperative α-blocker.

## Acknowledgements

We thank Lin-lin Jiang (Medical school of Chinese PLA, Beijing, China.), Xiao-fei Ye (Naval Medical University, Shanghai, China) and John Brunstein (Segra International Corp. Richmond, BC Canada) for they kind help during the study implementation and manuscript writing.

